# Unevolved proteins from modern and prebiotic amino acids manifest distinct structural profiles

**DOI:** 10.1101/2021.08.29.458031

**Authors:** Vyacheslav Tretyachenko, Jiří Vymětal, Tereza Neuwirthová, Jiří Vondrášek, Kosuke Fujishima, Klára Hlouchová

## Abstract

Natural proteins represent numerous but tiny structure/function islands in a vast ocean of possible protein sequences, most of which has not been explored by either biological evolution or research. Recent studies have suggested this uncharted sequence space possesses surprisingly high structural propensity, but development of an understanding of this phenomenon has been awaiting a systematic high-throughput approach.

Here, we designed, prepared, and characterized two combinatorial protein libraries consisting of randomized proteins, each 105 residues in length. The first library constructed proteins from the entire canonical alphabet of 20 amino acids. The second library used a subset of only 10 residues (A,S,D,G,L,I,P,T,E,V) that represent a consensus view of plausibly available amino acids through prebiotic chemistry. Our study shows that compact conformations resistant to proteolysis are (i) abundant (up to 40%) in random sequence space, (ii) independent of general Hsp70 chaperone system activity, and (iii) not granted solely by “late” and complex amino acid additions. The Hsp70 chaperone system effectively increases solubility and refoldability of the canonical alphabet but has only a minor impact on the “early” library. The early alphabet proteins are inherently more soluble and refoldable, possibly assisted by the cell-like environment in which these assays were performed.

Our work indicates that both early and modern amino acids are predisposed to supporting protein structure (either in forms of oligomers or globular/molten globule structures) and that protein structure may not be a unique outcome of evolution.

## Introduction

Today’s biological systems are anchored in the universal genetic coding apparatus, relying on coded amino acids that were likely selected in the first 10-15% of Earth’s history ^1^. While sources of prebiotic organic material provided a broad selection of amino acids, only about half of the canonical amino acids were detected in this pool ^2^. There is substantial evidence that this set formed an early version of the genetic code and that the “late” amino acids were recruited only after an early metabolism was in existence. The boundary between these two sets is blurry. However, large meta-analyses of these studies agree that “early”, i.e. the smaller and less complex amino acids (Gly, Ala, Asp, Glu, Val, Ser, Ile, Leu, Pro, Thr) were a fixture in the genetic code before its evolution ^3,4^.

The factors that drove the selection of 20 coded amino acids remain puzzling. Solubility, ease of biosynthesis, un/reactivity with tRNA, and potential peptide product stability seem to explain some selective “choices” but not others ^5,6^. Most recently, analysis of the *set* of amino acids revealed that the canonical alphabet shows an unusually good repertoire of the chemical property space when compared to plausible alternatives ^7,8^. Such studies lead to speculations that similar amino acid selection would be expected on other Earth-like planets ^5,8,9^.

In extant proteins, a significant part of the “late” amino acids (Arg, Lys, His, Cys, Trp and Tyr) belong to the essential catalytic residues, i.e. they are associated with catalysis in almost all of the enzyme classes ^10^. At the same time, the putatively early amino acids have been related to protein disorder and lack of 3D structure ^11^. However, sparse sampling of random sequences composed of early amino acids suggests that such proteins have a higher solubility than the full canonical alphabet ^12,13^. Moreover, computational and experimental mutational studies removing or reducing the late amino acids in selected proteins imply that the early amino acids comprise a non-zero folding potential ^14–18^. If prone to tertiary structure formation, it has been hypothesized that the early alphabet could more probably form molten globules rather than tightly packed structures, mainly due to the lack of aromatic and positively charged amino acids. According to this hypothesis, the addition of late amino acids would be required to increase protein stability and catalytic activity ^11,17,19^. Interestingly, it was shown that while positively charged amino acids are more compatible with protein folding, they also promote protein aggregation if their position within the sequence is not optimized or assisted by molecular chaperones. Thus it was hypothesized that chaperone emergence coincided with the incorporation of basic residues into the amino acid alphabet leading to an increase in the plasticity of natural folding space ^20^.

To assess the intrinsic structural and functional properties of the full amino acid alphabet, semi high-throughput studies using combinatorial sequence libraries have been performed previously ^21–25^. Surprisingly, secondary structure occurrence in random sequence libraries has been recorded with similar frequency as in biological proteins, while folding (or more precisely, occurrence of collapsed conformations) has been reported in up to 20% of tested proteins ^21,24,25^. However, more systematic and high-throughput screening is still necessary to confirm these observations. Moreover, it remains unclear how much these properties are a result of the full alphabet fine-tuning, whether structured molecules emerge spontaneously and independently in the canonical amino acid sequence space, and whether the early amino acids could provide similar structural traits.

To fill this knowledge gap, we characterized libraries of 10^12^ randomized protein sequences from the full and early amino acid alphabets to assess their collective biochemical characteristics. Our approach takes advantage of combinatorial samples to address the statistically inaccessible characterization of random sequence space by low throughput single-protein studies. While the bioinformatic prediction revealed similar secondary structure potential in both libraries and lower aggregation propensity of the full alphabet, the early alphabet is significantly more soluble and refoldable under cell-like experimental conditions. The full alphabet sequences were found to interact with molecular chaperones that can compensate for their otherwise poor solubility. Up to ∼40% of both library samples represent proteolysis resistant species. The results therefore agree with previous sparse sampling observations, and in addition, the folding frequency and inducibility of some properties in a cell-like environment are systematically mapped. Moreover, this study provides a unique synthetic biology pipeline that could be used to survey properties of any other protein alphabets associated with different biological phenomena of interest.

## Results

### Library expression and quality control

The combinatorial protein libraries studied in this work consisted of 105 amino acid long proteins with an 84 amino acid long variable parts, FLAG/HIS tag sequences on N’/C’ ends, and a thrombin cleavage site in the middle of the protein construct (Supporting Fig. S1). The variable region was designed by the CoLiDe algorithm and consisted of a specific set of degenerate codons in order to match the natural canonical (full alphabet, 20F) and the prebiotically plausible (A,S,D,G,L,I,P,T,E,V; early alphabet, 10E) amino acid distributions (Supporting Table S1) ^26^. The CoLiDe algorithm was chosen on the basis of its suitability for construction of vast combinatorial libraries. In comparison to alternative degenerate codon design tools it was specifically optimized for long variable protein libraries design rather than libraries suitable for site-specific mutagenesis investigations of protein variants. The design method consists in a selection of such degenerate codons which upon their combination in a degenerate DNA template produce a protein-coding library with the desired mean amino acid distribution. Although characteristics of different degenerate codons may produce a sequence-biased sample (each degenerate position will yield only a subset of the designed amino acid alphabet), this study aims to investigate effects of amino acid composition effects on random protein behaviour rather than sequence determinants of protein folding. The amino acid ratios for both libraries corresponded to natural amino acid distribution from the UniProt database ^27^. The libraries were assembled from two overlapping oligonucleotides, transcribed into their corresponding mRNA, and translated using an in vitro translation system (Supporting Fig. S2). In order to verify the designed library variability and amino acid distribution, we sequenced the assembled degenerate oligonucleotide DNA library and performed a mass spectrometric analysis of the purified library protein product. The root mean squared error (RMSE) from the target amino acid distribution was ∼0.06 in both libraries 20F and 10E (Supporting Table S2, Supporting Fig. S3). The variability analysis of the sequenced library showed that 96% of sequences were unique; no significant sequence enrichment was observed (Fig. 1, Supporting Table S3). Due to synthesis errors, STOP codons were introduced into 12% of the library sequences. The rates of misincorporation of undesired amino acids into library 10E did not exceed 1% and maximum deviation on single amino acid occurrence was 30% from the designed frequency (Supporting Table S2). The variability of the purified protein product was validated by MALDI-TOF mass spectrometry; the mean and spread of the experimental spectra closely matching the predicted distributions (Supporting Fig. S4).

**Figure 1.**
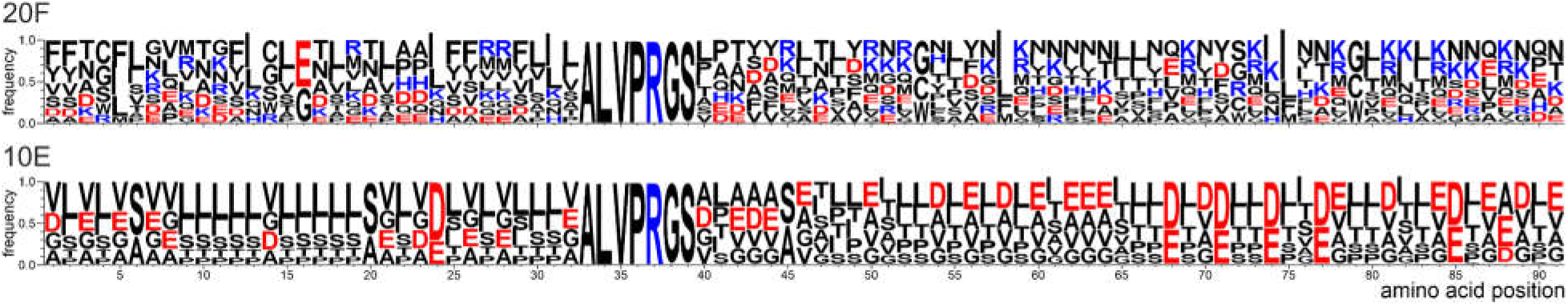
Sequence logo representation of full (top) and early (bottom) alphabet libraries variability constructed from the corresponding sequenced DNA templates. Sizes of the letters represent frequencies of specific amino acids per each position in the set of translated sequenced templates. Proteins coded by degenerate DNA templates can be represented by a linear combination of all residues with each amino acid occurring with its distinct frequency.

### Secondary structure and aggregation propensity predictions

Sequences of both 20F and 10E libraries acquired by high throughput sequencing were analyzed by a consensus protein secondary structure prediction ^28^. 200,000 sequences were analyzed from each library. Interestingly, despite the different amino acid distributions, comparable alpha helix and beta sheet forming tendencies were reported in both libraries with only a slight increase in alpha helix content in the 20F library (33 % vs. 30% in 10E) (Fig. 2A). The overall alpha helix and beta sheet content correlate well among the individual predictors used for both studied libraries, which is not necessarily the case for other alternative and more artificial alphabets (unpublished observation). The prediction of aggregation propensity of the same set of sequences indicated a significantly higher aggregation tendency of 10E library proteins in comparison to 20F library proteins (Fig. 2B).

**Figure 2.**
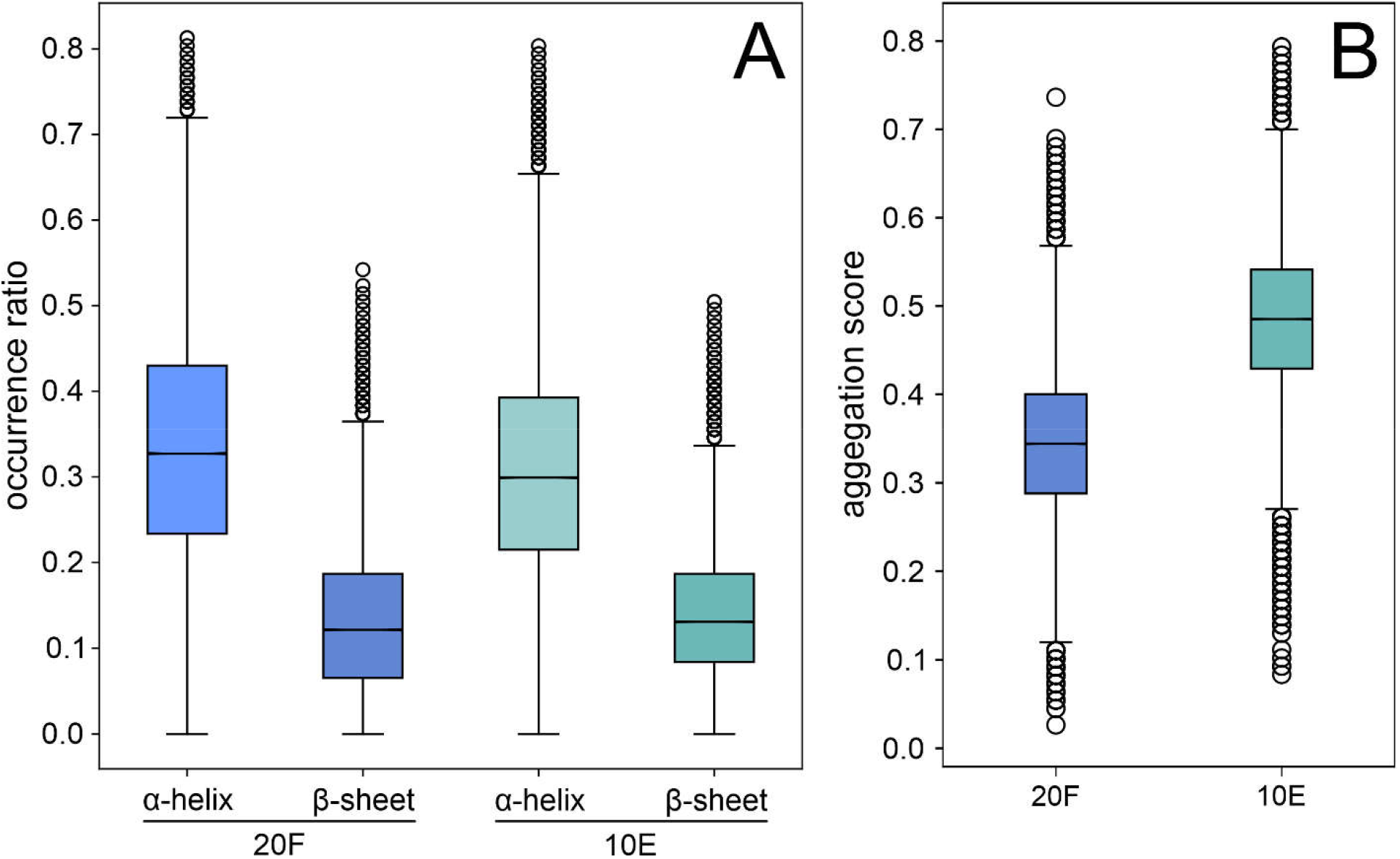
Bioinformatic prediction of alpha helix and beta sheet content (A) and aggregation propensity (B) of a sample of 200,000 sequences acquired by high throughput sequencing of the early (green) and full (blue) alphabet library DNA templates. Aggregation score is defined as the ratio of predicted aggregation-prone residues per sequence. The box extends from first quartile to third quartile with a line in the middle representing the median. The whiskers extend from the box by 1.5x the inter-quartile range. The points above and under the boxplot represent sequences whose predictions fall outside the 1.5x the inter-quartile range from the box edges

### Expression and solubility analysis in the absence and presence of the DnaK chaperone system

To systematically assess the expression profiles of the libraries, a quantitative western blot analysis was performed with the library products expressed at different temperatures (25, 30 and 37 °C) and with/without DnaK/DnaJ/GrpE chaperone system supplementation (further referred as to DnaK). The analysis was carried out in triplicate, and western blot signals of both total expression and soluble fractions were quantified with ImageJ ^29^. For both 20F and 10E libraries, the expression yields grow with increasing temperature, with the overall yield being mildly lower in the chaperone supplemented reactions at 37 °C (Fig. 3). In the case of the 20F library, the solubility of the library is relatively poor but is significantly improved by chaperone supplementation. While in the chaperone supplemented reaction the soluble fraction grew with expression temperature proportionally with the total expression, in the chaperone absent condition, the soluble fraction yields did not significantly change with the transition from 30 to 37 °C. On the other hand, chaperone supplementation did not have a significant effect on the 10E library expression or solubility (Fig. 3).

**Figure 3.**
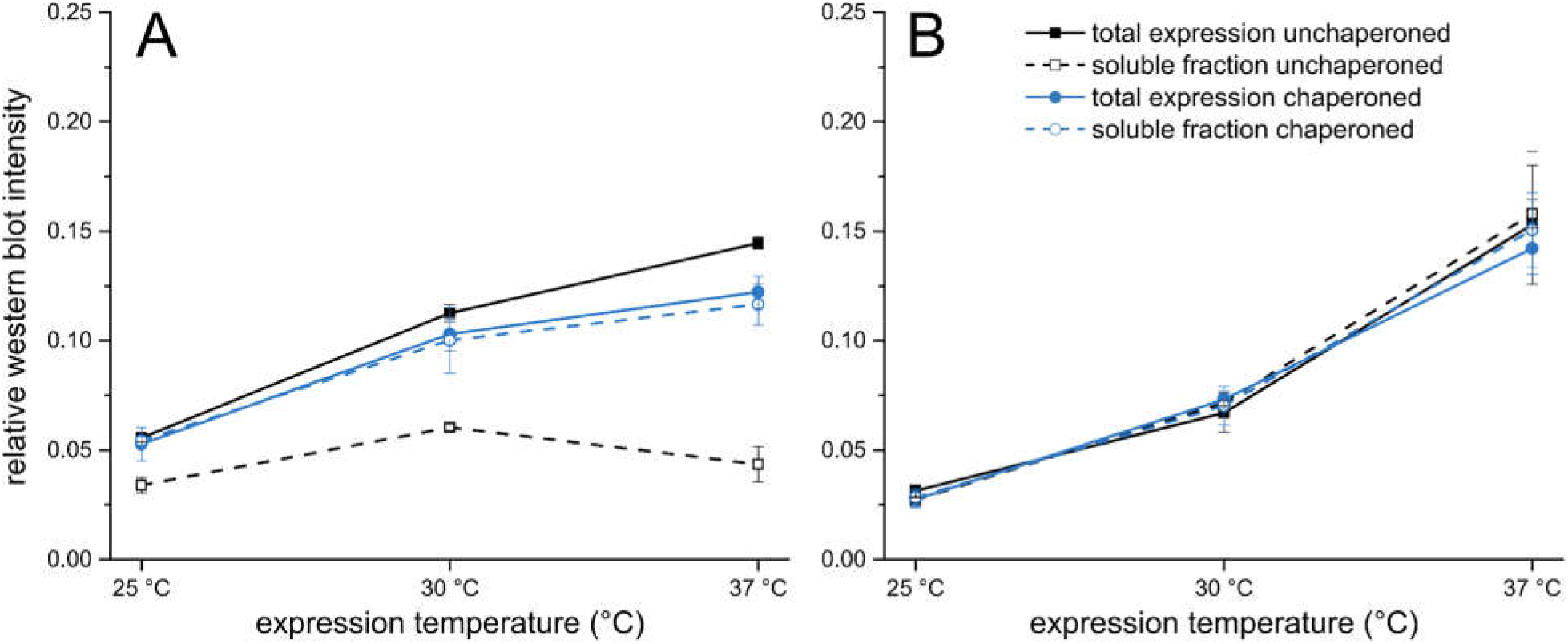
A summary of expression and solubility analysis of the full (A) and early (B) alphabet libraries at three different temperatures. Total expression (solid line) and soluble fraction (dashed line) were compared in chaperoned (blue line) and unchaperoned (black line) conditions. For original data see Supporting Fig. S5/S6 and Supporting Table S4.

### Assessment of proteolytic resistance

The structural potential of random protein libraries was assessed by proteolysis. The digestion assessment was performed in triplicate by Lon and thrombin proteases in co-translational and post-translational conditions, respectively (Fig. 4). The Lon protease is a part of the *E. coli* protein misfolding system and is known to specifically digest unfolded proteins in exposed hydrophobic regions ^30^. Here we adapted a previously published protocol on single protein structure assessment for combinatorial library characterization ^31^. The method is used to separate and quantify distinct protease sensitive parts of the library within both the soluble and insoluble fractions of the expressed libraries. The thrombin protease assay was adapted from the study of Chiarabelli et al, wherein the structure occurrence is derived from the cleaved/uncleaved ratio of proteins with an engineered thrombin cleavage site situated in the middle of the sequence ^21^. The unstructured proteins are expected to be quickly degraded on the exposed cleavage site.

**Figure 4.**
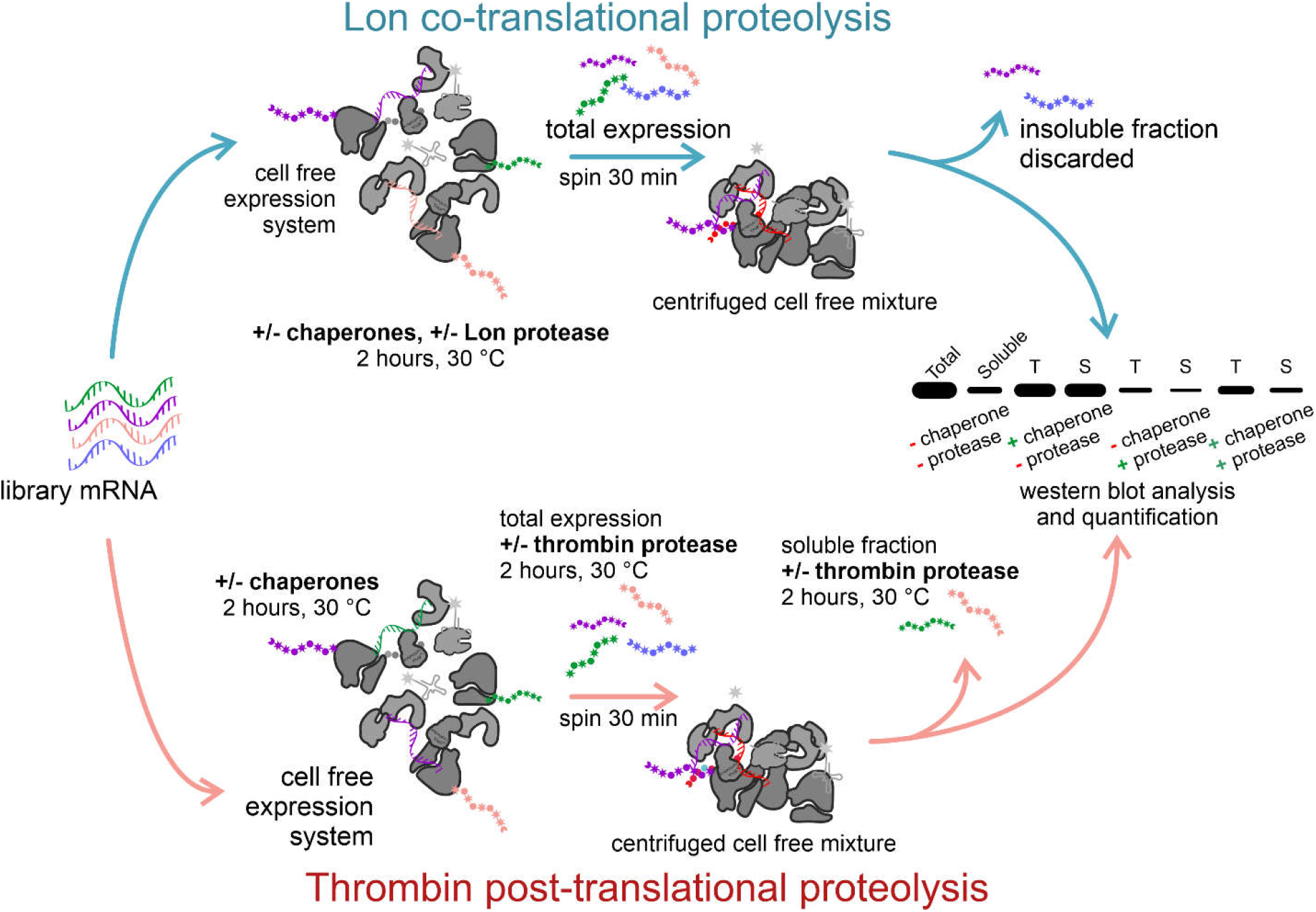
Scheme of the proteolytic resistance experimental pipeline. In the co-translational proteolytic assay (top) the Lon protease is present during the cell-free expression; in the post-translational proteolytic assay (bottom) thrombin protease is added to the separated total and soluble fractions of the expressed library after translation is quenched by addition of puromycin

While co-translational Lon protease assay represents real-time analysis of protein folding kinetics, thrombin protease digestion aims for an indirect final folding assessment via proteolysis on accessible or buried cleavage sites. Both assays target different stages of protein folding pathways and bring distinct insights into the overall random protein folding behaviour.

According to the 20F library analysis, the soluble/undegradable structured proteins represent ∼30-35% of the total product (Fig. 5A). Upon addition of the DnaK chaperone, most of the library solubilizes, but the protease resistant content does not increase significantly and occupies ∼40-50% of the total product. In comparison, chaperone addition does not have an impact on the solubility or protease resistance of the 10E library (Fig. 5B). Interestingly, the protease resistant content (soluble undegradable) in the 10E library is similar as in the 20F library after the addition of chaperones.

**Figure 5.**
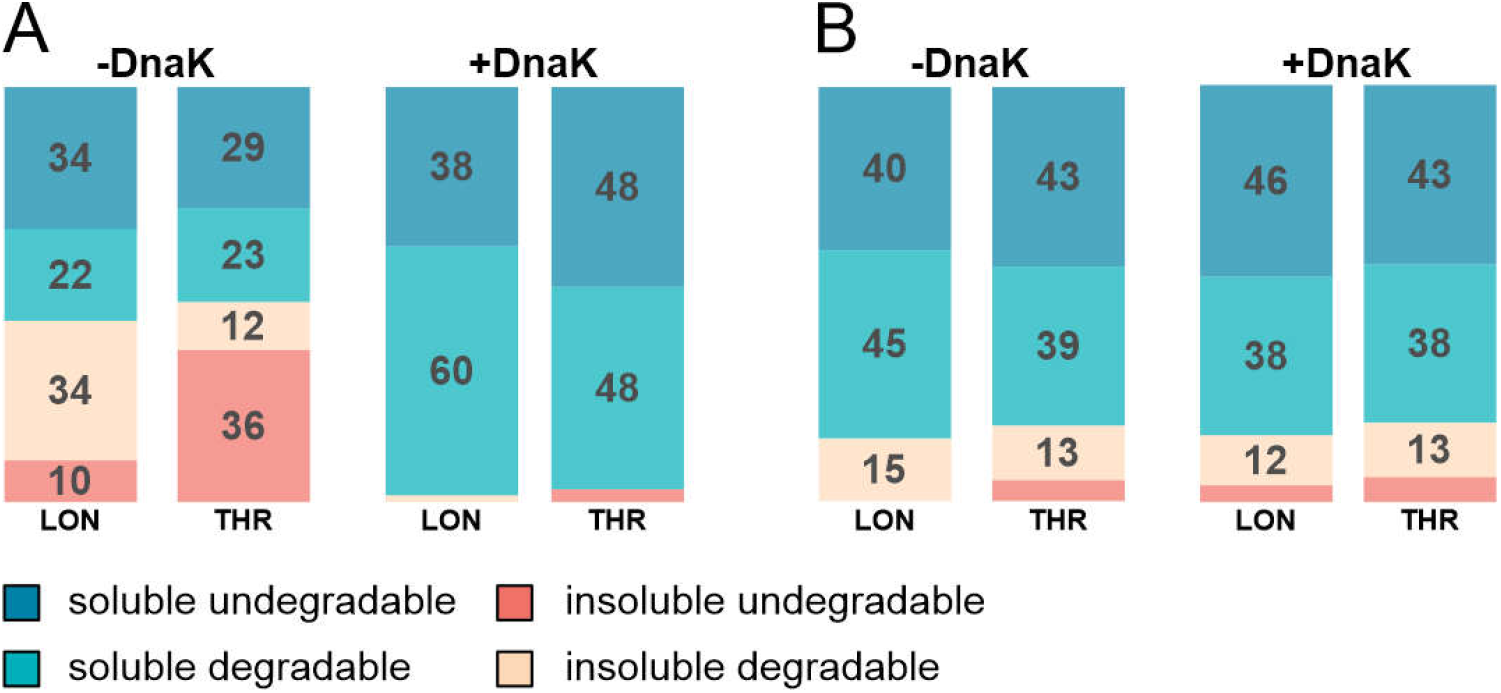
An integrated solubility/proteolysis resistance analysis of the full (A) and early (B) alphabet libraries. Libraries were expressed either in the absence (left double column) or presence (right double column) of the DnaK chaperone system. Proteolysis was performed by protease Lon (left columns) in a co-translational regime or by thrombin protease (right columns) in a post-translational mode. Values in the boxes represent the percentage ratios of the total expressed library per fraction. For original data see Supporting Fig. S7/S8/S9/S10 and Supporting Table S5/S6

### Protein heat shock refoldability characterization

Following expression, solubility, and protease resistance assessment, we analyzed the temperature sensitivity of the 20F and 10E proteins. The libraries expressed with and without chaperone supplementation were subjected to 15 minutes/42 °C heat shock. The aggregated fraction was removed by centrifugation, and the soluble fraction was compared with and without thrombin treatment (Fig. 6).

**Figure 6.**
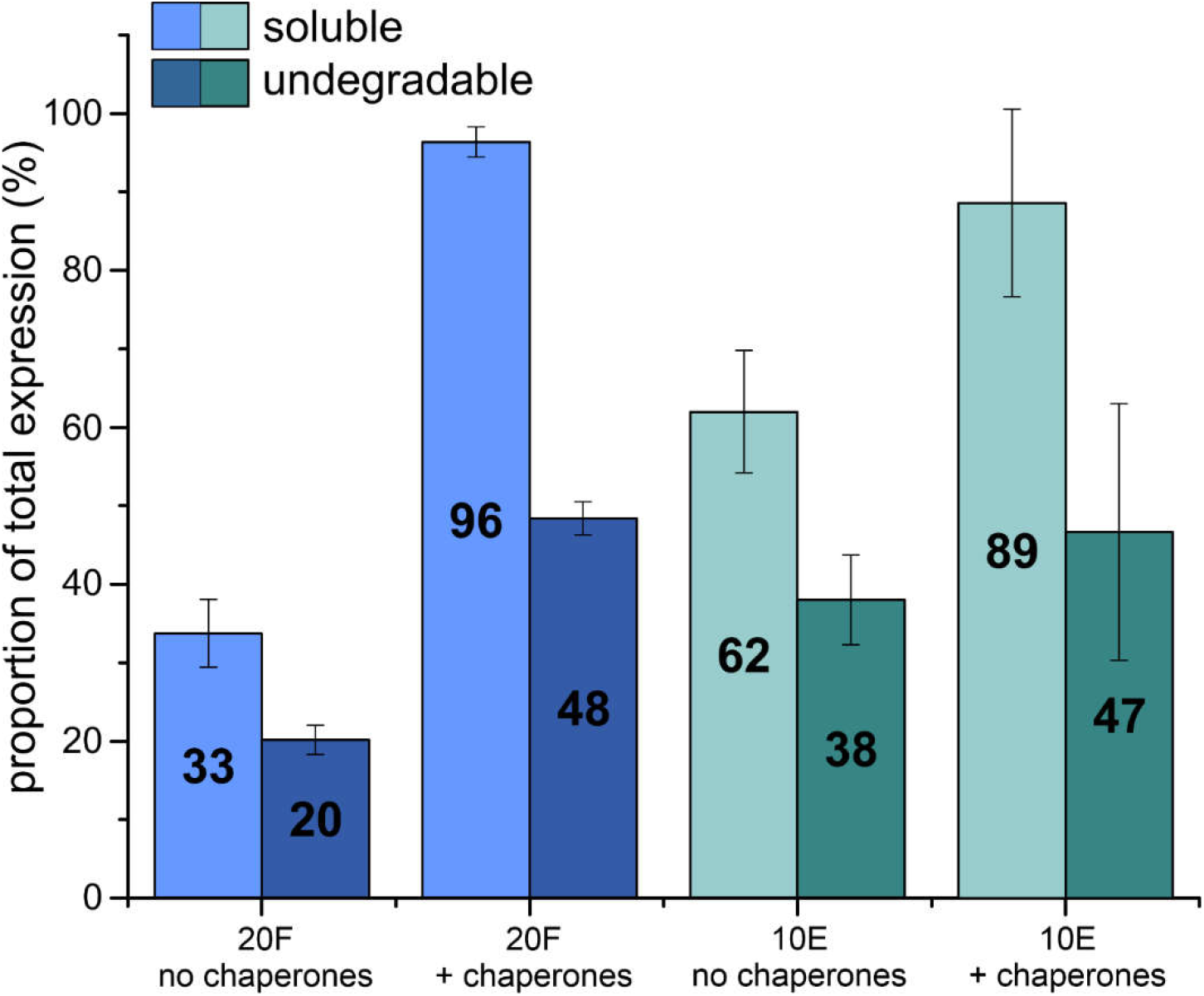
Thermostability analysis showing soluble proportions (light blue and green) of the total expression of the full and early alphabet libraries after a heat shock (42 °C / 15 min) treatment and their respective thrombin resistant proportions (dark blue and green) of the total expression in unchaperoned and chaperoned conditions. Numbers in the bars represent the percentage fraction of the total expressed library. For original data see Supporting Fig. S9/S10 and Supporting Table S6.

The 10E library is intrinsically more soluble than 20F (∼60 vs ∼30% of the libraries remain soluble after heat shock, respectively) while the DnaK chaperone system induces higher post-heat shock solubility in both libraries. The protease resistant fraction of the soluble part of the libraries remains the same (∼40%) as before heat shock treatment with the exception of the unchaperoned 20F library, which demonstrates a slight decrease in both the soluble and degradation resistant fractions (Fig. 6).

## Discussion

In this study, a high-throughput systematic approach was used to experimentally analyze the structural properties of the vast protein sequence space. Random sequences have been proposed as proxies for both precursors of *de novo* emerged proteins in current evolution as well as (ii) sources of peptide/protein birth at the earliest stages of life preceding templated proteosynthesis ^32,33^. However, the structural properties of random sequences have so far remained uncomprehended, while a few recent bioinformatic and coarse-grained studies have pointed to their surprising properties, such as high secondary structure propensity and *in vivo* tolerance ^24,25,34^. Here, two combinatorial protein libraries encompassing upto 10^12^ individual sequences from two distinct alphabets (representing hypothetical stages of genetic code evolution) have been characterized.

### Solubility of the natural alphabet random proteins can be induced by chaperones

The first “full” alphabet library is based on the amino acid composition of the Uniprot database representing the properties of today’s proteomes. It has previously been shown that similar constructs have limited solubility but a similar secondary structure potential to biological proteins ^12,13,25^. Our study confirms these results, and in addition, we specify that 20-50% of the overall diverse library appears in the soluble fraction in the 30-37 °C temperature range. No significant aggregation was observed upon the library expression in 25 °C. While previous studies of similar construct size evaluated the solubility of individual proteins that were overexpressed (many of them with partial solubility) in different *E. coli* strains and under different conditions, our library was expressed using a reconstituted cell-free protein synthesis (CFPS) system, and its large diversity (contrasting with overexpression of individual proteins) was confirmed by MALDI. Therefore, we cannot make a direct comparison to previous studies of individual proteins but rather report the “fingerprint” properties of the full alphabet domain-size proteins.

Interestingly, this library of unevolved sequences was observed to interact productively with the natural molecular chaperone system DnaK/DnaJ/GrpE which was used to supplement the CFPS system in another experiment. This interaction caused almost total solubilization of the otherwise insoluble proteins over the studied temperature range. While the solubility traits may be quite different for much shorter polymer lengths, our previous study showed that random domain-size sequences cope with significant aggregation, especially if they are rich in secondary structure content ^25^. To characterize the library structural potential without introducing potential bias, we used an *in situ* double proteolysis experiment adapting two previously reported approaches ^21,31^. The experiment combined co-translational proteolysis by disorder-specific Lon protease and a post-translational cleavage by thrombin designed to cut the potentially exposed cleavage site engineered in the center of random proteins. Besides the increased robustness of the structure content estimation, such a combined approach provides unique insight into the library translation dynamics. The double proteolysis experiment revealed that ∼30-35% of library 20F proteins are protease resistant. Upon the addition of chaperones (which solubilizes the library as described above), the ratio of protease resistant species rose only mildly to ∼40-50%. The more prevalent protease sensitive nature of the full alphabet library echoes the reported nature of *de novo* proteins, i.e. proteins that emerge in current biology from previously non-coding DNA (summarized in ^35^).

Overall, these results show that while inherent protein solubility is limited in random sequence space made of the natural alphabet, it can be induced significantly by the activity of molecular chaperones. At the same time, the DnaK chaperone system has only a minor effect on the level of protease resistance, suggesting that the majority of the potentially solubilized sequences are devoid of higher structure arrangements. In comparison, the same folding assessment of 76 randomly selected *E*.*coli* proteins by Niwa et al. were Lon-resistant in their soluble fraction, suggesting a high level of folding optimization of biological vs. random proteins ^31^. These results are in agreement with earlier studies of the Hecht group which pointed out that even though structured arrangements are achievable within random sequence space, most of its representants appear to be in relaxed molten globule states ^36,37^. Nevertheless, the ∼40% natural abundance of soluble and yet protease-resistant sequences in unevolved sequence space may be surprising in light of earlier hypotheses and even exceeds the estimates of folding frequency reported by previous coarse-grained studies ^21,38^. However, major differences in the experimental setups (cell-free vs cell-based expression, low-level vs overexpression, high- vs. low-throughput methodology, library design and sequence length) prevent the possibility of direct comparisons among these studies. A direct comparison of the full library properties can however be made with another library of proteins studied here under the same experimental conditions.

### Protease resistance is comparable in proteins from the full canonical alphabet and its early subset, unaffected by chaperones

A second “early” alphabet library was constructed from a 10 amino acid subset of the full alphabet which was proposed to constitute an earlier version of the genetic code and be reflected in the composition of early proteins ^3^. We emphasize that with this study, we do not try to establish that there was necessarily a time in life’s evolution during which domain-size proteins were composed entirely of this amino acid subset. Our analysis rather deals with the inherent physico-chemical properties of such an alphabet, were it to form or dominate protein-like structures. We also acknowledge that the earliest stages of peptide/protein formation (preceding templated proteosynthesis and perhaps also its early less specific versions) probably utilized a plethora of prebiotically plausible amino acids or similar chemical entities, but inclusion of such non-canonical amino acids in the studied alphabets is beyond the scope of this study ^1,39,40^.

Although the overall secondary structure propensity of the early alphabet is comparable to the full alphabet, according to the bioinformatic prediction, the occurrence of alpha-helix is slightly (∼3%) lower. While these differences are statistically borderline, they may have interesting implications for the evolution of protein structural properties. Brack and Orgel proposed that beta-sheet structures were prebiotically significant, and the later significance of alpha-helices in protein folds was also recently implied by the structural analysis of ribosomal protein content, showing that the most ancient protein-protein fragments of this molecular fossil are mostly disordered and of beta-sheet formation ^41–43^. Despite the similar secondary structure propensities of the full and early alphabets, the 10E library proteins are significantly more soluble (∼90%) upon expression. They retain similar solubilities in chaperoned/unchaperoned conditions unlike the 20F library proteins. This observation supports the previously stated hypothesis of chaperone co-evolution with the incorporation of the first positively charged amino acids into the early amino acid alphabets ^20^. That way proteins composed of the full alphabet would be kept in solution despite their higher inherent aggregation propensities.

The significantly higher solubility of the 10E library proteins (and similar protein compositions) is in agreement with previous studies ^12,13^. This phenomenon could be related to the lower complexity of 10E library proteins resulting from the limited amino acid alphabet. While 20F proteins represent a highly variable sample of protein folding space with many opportunities for aggregation initiation, the 10E proteins display a narrower subspace with much more uniform sequence and physicochemical characteristic distributions. In addition, their overall negative charge and absence of positively charged/aromatic amino acids are factors which were previously shown to suppress both nonspecific aggregations as well as independent protein folding formation ^20,44^. At the same time though, the 10E alphabet contains a significant proportion of hydrophobic amino acids. Using the ProA bioinformatic predictor of protein aggregations, the 10E library would be expected to be intrinsically less soluble, contradicting our observations as well as previous empirical observations. However, contrasting with the intrinsic behavior of the protein alone, our assays (and previous experimental assays) were performed in a cell-like environment, rich in different salts and other small molecules/cofactors. This might have implications discussed further below.

Interestingly, the 10E library also displays a significant protease resistant behaviour. In the absence of chaperones, the ratio of the protease resistant fraction is 40-50% in both the co- and post-translational digestion assay, i.e. similar to the 20F protease resistant fraction when supplemented with chaperones.

Such a high level of protease resistance within the 10E library is non-intuitive and unexpected purely from its amino acid composition. However, several folders have been recently identified from the same or similar protein composition in experiments reducing extant protein compositions ^15,16,18,45,46^. Where characterized in more detail, assistance of salts, metal ions, or cofactor binding were found to explain the folding properties ^15,18,45,47^. In addition, Despotovic et al. recently confirmed that folded conformations of a highly acidic 60-residue protein can be induced by positively charged counterions, in case of Mg^2+^ the reported concentration corresponding roughly to its concentration in the CFPS reaction (∼10mM) ^48^. In parallel, the Hecht group reported that binding of metal ions (with high nanomolar to low micromolar affinity) is a surprisingly frequent property of unevolved sequences and therefore does not require much sequence optimization ^49^. These studies allow us to speculate that the high protease resistance of the 10E alphabet could result from the folding assistance in cation/cofactor-rich environment, where the lack of hydrophobic and electrostatic interactions is compensated by these chemical entities. Alternatively or concurrently, the library solubility and protease resistance could be partly explained by tertiary structure formation induced by oligomerization as previously hypothesized by Yadid et al. in a study using 100 amino acid long fragments (albeit from different amino acid compositions) ^50^. Our study presented here cannot unambiguously differentiate between these two possible scenarios or their combination as the highly variable library sample of a limited amount prevents more sophisticated physico-chemical analyses that could be used to address these phenomena in follow-up studies.

### Early alphabet proteins are inherently more temperature resistant in a cell-like milieu

One of the notable assumed characteristics of the early prebiotic Earth is the elevated temperature of the environment ^51^. The temperature-induced aggregation propensity of random protein libraries was investigated by their exposure to a mild 15-min heat shock at 42 °C. Interestingly, the quantity of soluble fractions in proteins without chaperones were approximately two times greater in the early alphabet library (∼30% vs ∼60% for 20F and 10E libraries, respectively) which might indicate a natural tendency to withstand elevated temperature. On the other hand, addition of chaperones decrease aggregation tendencies of both 20F and 10E libraries up to almost full solubility upon heat shock treatment. This observation confirms our previous conclusions about the strong dependence of the canonical amino acid alphabet proteins on chaperone activity and extends it to aggregation prevention of the early amino acid alphabet proteins. Additionally, the fraction of protease resistant proteins remains unchanged (∼40%) upon heat shock for both libraries, suggesting that the proteins destabilized by elevated temperature belong to the unstructured category.

While most of the above referenced studies reducing the composition of extant proteins towards the early set of amino acids did not observe an increase in their temperature resistance ^15,16,18,45,47^, we are here concerned with a comparison of unevolved sequences from the two amino acid repertoires and their inherent properties.

### Concluding remarks

In summary, while our study confirms some of the previously reported properties of the random sequences space (such as its surprisingly high secondary structure potential and relative ease of expression), we expand on this knowledge using a systematic high-throughput approach using diverse combinatorial libraries composed of two different alphabets. Escaping the restraints of sparse sampling, our study maps protease resistance, solubility, and temperature resistance in random sequences composed of the natural vs. the early evolutionary canonical alphabets. Along with the advantages of the high-throughput approach to directly compare the two protein alphabets, the methodology applied in our study is inevitably limited by the nature of the library samples. Although the sample sizes in our experimental study are still minuscule in comparison to the vast potential random sequence space, this work presents a qualitative view on structure forming potential within the unevolved protein domain. Protease resistance serves as a rather low resolution technique to study the overall structural propensities rather than specific tertiary structure arrangements. Future studies would be needed to address detailed structural and functional properties of purified proteins that can be selected from the diverse libraries. The analyses presented here were performed in a cell-like environment (rich in salts and cofactors) that may better represent protein formation conditions during both the origins of life and in extant biology. Under such conditions, the early alphabet sequences (i) are inherently more soluble and (ii) remain in solution when unfolded. These properties are partially achievable to full amino acid alphabet proteins through interaction with molecular chaperones which suggests a compelling argument for protein chaperone activity evolution. Interestingly, our study reports that both alphabets frequently give rise to proteolysis resistant soluble structures, occupying up to ∼40% of all sequences. Because the intrinsic properties of the prebiotically plausible amino acids do not imply such properties, we hypothesize that the protein solubility and folding within this library are enabled by the cell-like milieu, assisted by salts, metal cations, and cofactors. Follow up studies are suggested to further explore these findings as our initial proteolytic structure assessment does not allow for differentiation of various flavors of protein structure such as oligomeric assemblies, molten globular or stable hydrophobic globular arrangements.

## Methods

### Design of libraries from early and full amino acid alphabet

Two 105 amino acid long random sequence libraries were designed using the CoLiDe algorithm for combinatorial library design ^26^ and the amino acid ratios listed in Supporting Table S1. The randomized part of the libraries consisted of 84 amino acids; the remainder is attributed to the FLAG affinity purification site on the N-end of the construct, the hexahistidine tag on the C-end, and the and thrombin protease recognition site (ALV**PRG**S) in the middle of the construct (Supplementary Figure S1).

### Bioinformatic analysis of secondary structure potential

Prediction of secondary structure potential of the studied libraries was performed by a consensus predictor as described previously ^28^. It combines outputs of the spider3, psipred, predator, jnet, simpa, and GOR IV secondary structure predictors ^52–57^. None of the predictors were allowed to use homology information that might prevent high-throughput processing of protein sequences. The final assignment of secondary structure followed the most frequently predicted secondary structure element at each amino acid position. Protein aggregation was predicted by the ProA algorithm in a protein prediction mode ^58^. The protein sequences and prediction results are available at the OSF platform website (https://osf.io/4e9s2/).

### Preparation of experimental libraries

20F and 10E DNA libraries were synthesized commercially as two overlapping degenerate oligonucleotides (see Supplementary information for the sequences) that were designed by the CoLiDe algorithm to follow the natural canonical (full alphabet, 20F) and prebiotically plausible (A,S,D,G,L,I,P,T,E,V; early alphabet, 10E) amino acid distributions (Supporting Table S1). The overlapping oligonucleotides were annealed and extended by Klenow fragment to form double-stranded DNA (dsDNA). Annealing was performed by heating the complementary oligonucleotide mixture (48 μl total reaction volume, 2 μM final concentration of each) in NEB2 buffer provided with 200 μM dNTPs to 90 ºC for 2 minutes and cooling down to 32 ºC with a 1 ºC/min temperature gradient. The Klenow extension was performed by Klenow polymerase (NEB): 10 U of Klenow polymerase was added to annealed oligonucleotides, incubated for 5 minutes at 25 °C, 37°C for 1 hour (polymerization step), and 50 °C for 15 minutes (inactivation step). Final dsDNA libraries were further column purified using the DNA Clean and Concentrator kit (Zymo Research), and the product was quantified by Nanodrop 2000c (Thermo Scientific). In the following transcription, 1 μg of DNA library was used as a template for mRNA synthesis by HiScribe T7 kit (NEB). The product was purified by NH4Ac precipitation and dissolved in RNAse-free water to a final concentration of 3 μg/ul.

The library DNA was analyzed by high throughput sequencing on Illumina MiSeq. The libraries for next generation sequencing (NGS) were prepared from 100 ng DNA samples using the NEBNext Ultra II DNA Library Prep kit (New England Biolabs) with AMPure XP purification beads (Beckman Coulter). the length of the prepared library was determined by Agilent 2100 Bioanalyzer (Agilent Technologies) and quantified by Quantus Fluorometer (Promega). The sample was sequenced on a MiSeq Illumina platform using the Miseq Reagent Kit v2 500-cycles (2×250) in a paired-end mode. Raw data was processed with the Galaxy platform, and sequence analysis of assembled and filtered paired reads was performed with MatLab scripts developed at Heinis lab ^59,60^. The raw sequencing data is available at OSF platform website (https://osf.io/4e9s2/). the protein library was expressed using the PUREfrex 2.0 (GeneFrontier Corporation) recombinant in vitro translation system. The reaction was supplemented by 0.05 % (v/v) Triton X-100 and prepared according to manufacturer recommendations. The reaction was initiated by 3 μg of library mRNA. Expression followed for 2 hours at 25, 30, or 37 °C.

### Affinity purification of protein libraries

Expressed protein libraries were diluted 10x in binding buffer (50mM Tris, 150 mM NaCl, 0.05% (v/v) Triton X-100, pH 7.5) and incubated for 2 hours at 25 °C with 3 μl / 20 μl reaction of TALON affinity purification matrix. The immobilized library was washed three times with binding buffer and eluted by addition of 20 μl / 20 μl reaction of elution buffer (50mM Tris, 150 mM NaCl, 10mM EDTA, 0.05% (v/v) Triton X-100, pH 7.5).

### Solubility analysis of protein libraries

Cell free protein expression reactions were supplemented with 0.05 % Triton X-100, and protein libraries were expressed in different temperatures according to manufacturer recommendations. In order to analyze the quantity of total protein product, 10 μl of each reaction was quenched by addition of 40 μl of 300 μM puromycin in 50 mM Tris, 100 mM NaCl, 100 mM KCl, pH 7.5. Quenching proceeded for 30 minutes at 30 °C. Next, 5 μl of the quenched reaction mixture was taken for the following SDS-PAGE analysis of total library expression; the rest of the mixture was centrifuged for 30 minutes at 21 °C, and 5 μl of supernatant was taken for SDS-PAGE analysis of the soluble fraction of the library. Both fractions were analyzed by quantitative Western blotting (Sigma-Aldrich Monoclonal ANTI-FLAG® M2-Peroxidase (HRP) antibody, A8592) following the SDS-PAGE separation.

### Lon proteolytic assay of protein libraries

Lon protease was expressed and purified according to the previously published protocol ^31^. Cell free expression reactions were supplemented with 0.05 % Triton X-100; reactions were prepared according to manufacturer recommendations. Libraries were expressed in the presence or absence of the DnaK chaperone (K+/K-) and in the presence or absence of Lon protease (L+/L-). Chaperones were added to the final concentration of 5 μM DnaK, 1 μM DnaJ, 1 μM GrpE and Lon protease to 0.4 μM (hexamer)/reaction. Expression proceeded in 10 μl reaction volume for 2 hours at 30 °C and was quenched by 40 μl addition of 300 μM puromycin in 50 mM Tris, 100 mM NaCl, 100 mM KCl, pH 7.5. Quenching proceeded for 30 minutes at 30 °C. The sample preparation of total and soluble library fractions was identical to the solubility analysis experiment described above.

### Thrombin proteolytic assay of protein libraries

Cell free expression reactions were supplemented with 0.05 % Triton X-100; reactions were pre-pared according to manufacturer recommendations. Libraries were expressed in the presence or absence of the chaperone DnaK (K+/K-). Chaperones were added to the final concentration of 5 μM DnaK, 1 μM DnaJ, 1 μM GrpE μM. Expression proceeded in 10 μl reaction volume for 2 hours at 30 °C and was quenched by 40 μl addition of 300 μM puromycin in 50 mM Tris, 100 mM NaCl, 100 mM KCl, pH 7.5. Quenching proceeded for 30 minutes at 30 °C. Post-translational thrombin proteolysis was prepared as follows: 5 μl of quenched reaction was diluted 4x by 15 μl of 50 mM Tris, 100 mM NaCl, 100 mM KCl, pH 7.5; 0.15 U of thrombin (SigmaAldrich, USA) was added, and the total expressed library was digested for 2 hours at 30 °C. The soluble fraction of the library was prepared by centrifugation at 21 000 xg for 30 minutes at 21 °C, and 5 μl of supernatant was thrombin digested according to the same protocol. Cleaved samples of the total expressed and soluble libraries were analyzed by SDS-PAGE and Western blotting (Sigma-Al-drich Monoclonal ANTI-FLAG® M2-Peroxidase (HRP) antibody, A8592). The final quantification was performed on undigested fractions of the library proteins rather than on formation of cleavage frag-ments due to experimental errors in small fragments transfer and secondary cleavage of formed fragments of library 20F.

### Thermostability assay

Libraries expressed in 10 μl volume were processed as described above in the Lon proteolytic assay protocol. The Lon absent libraries were further analyzed for their thermostability in the presence and absence of chaperone. Processed reactions were incubated at 42 °C for 15 minutes and immediately centrifuged at 21 000 ×g for 30 minutes at 21 °C. The 5 μl supernatant fractions were subjected to thrombin proteolysis as described previously and analyzed by SDS-PAGE and quantitative western blotting.

### Quality control of purified protein libraries

For mass spectrometry, the purified protein library sample was resuspended in water. The spectrum was collected after addition of 2,5-dihydroxybezoic acid matrix substance (Merck) using an UltrafleXtremeTM MALDI-TOF/TOF mass spectrometer (Bruker Daltonics, Germany) in linear mode.

## Supporting Information

Supporting Table S1: Summary of amino acid and degenerate codon contents of the designed libraries

Supporting Table S2: Statistics on amino acid content of designed and experimentally validated libraries

Supporting Table S3: Statistics on library multiplicity and correctness of sequenced DNA templates

Supporting Table S4: Numerical output of western blot analysis of libraries solubility in different conditions

Supporting Table S5: Numerical output of western blot analysis of libraries LON proteolysis assay

Supporting Table S6: Numerical output of western blot analysis of libraries thrombin proteolysis assay

Supporting Fig. S1: Schematic representation of designed library template

Supporting Fig. S2: Agarose gel with DNA/RNA templates involved in library characterization

Supporting Fig. S3: Comparison of designed vs experimental amino acid rations within the combinatorial libraries

Supporting Fig. S4: MALDI-TOF MS spectrum of the purified combinatorial libraries overlapped with their theoretical mass distribution

Supporting Fig. S5,S7,S9: Western blot images used for quantification of libraries solubility and protease resistances

Supporting Fig. S6,S8,S10: Graphical representation of averaged signal intensities from western blot quantification of libraries solubilities and protease resistances

Supporting sequences: DNA templates of the expression cassettes used to express combinatorial libraries in cell-free expression system

## Acknowledgements

We are grateful to Prof. Hideki Taguchi and Prof. Tatsuya Niwa for kindly providing us with the expression plasmid of the Lon protease used in this study. We would also like to acknowledge Dan S. Tawfik^†^ and Valerio Guido Giacobelli for helpful discussions regarding this manuscript. In addition, we would like to thank Kateřina Nováková for her technical help with collecting MALDI spectra. This work was supported by the Czech Science Foundation (GAČR) grant number 17-10438Y and the Human Frontier Science Program grant HFSP-RGY0074/2019. K.F. is supported by ELSI - First Logic Astrobiology Donation Program.

## Author Contributions

VT and KH designed research; VT, JVy, TN and KF performed research; VT, JVo, KF and KH analyzed data; VT and KH wrote the paper.

