## Supplementary material for "Unevolved proteins from modern and prebiotic amino acids manifest distinct structural profiles"

### Supporting Figures

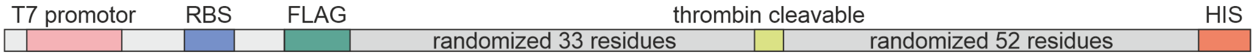


*Supporting Figure S1. The general scheme of the library expression cassette. Construct is provided with necessary sequences for in vitro transcription and translation, FLAG/HIS affinity purification tags on N/C ends of proteins and the thrombin cleavage site in the middle of the protein coding sequence*

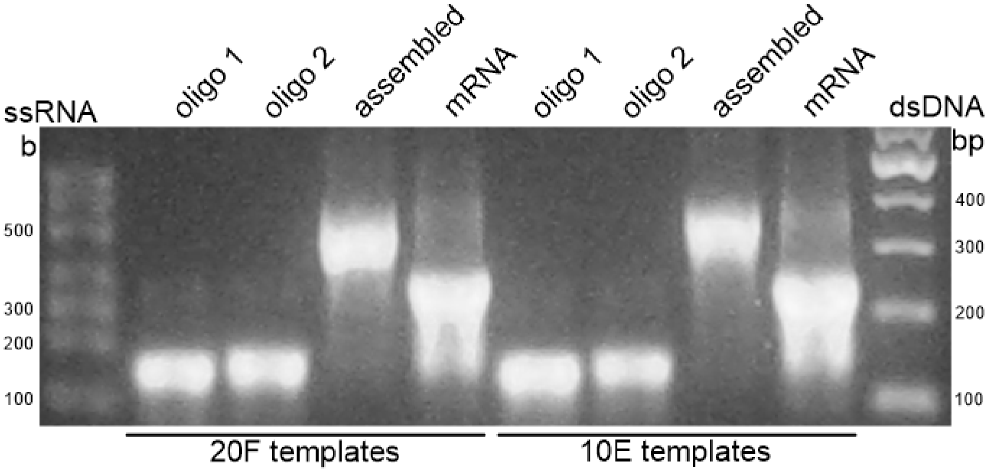


*Supporting Figure S2. Agarose gel representing degenerate ssDNA and assembled dsDNA library templates (A) and transcribed mRNA templates (B)*

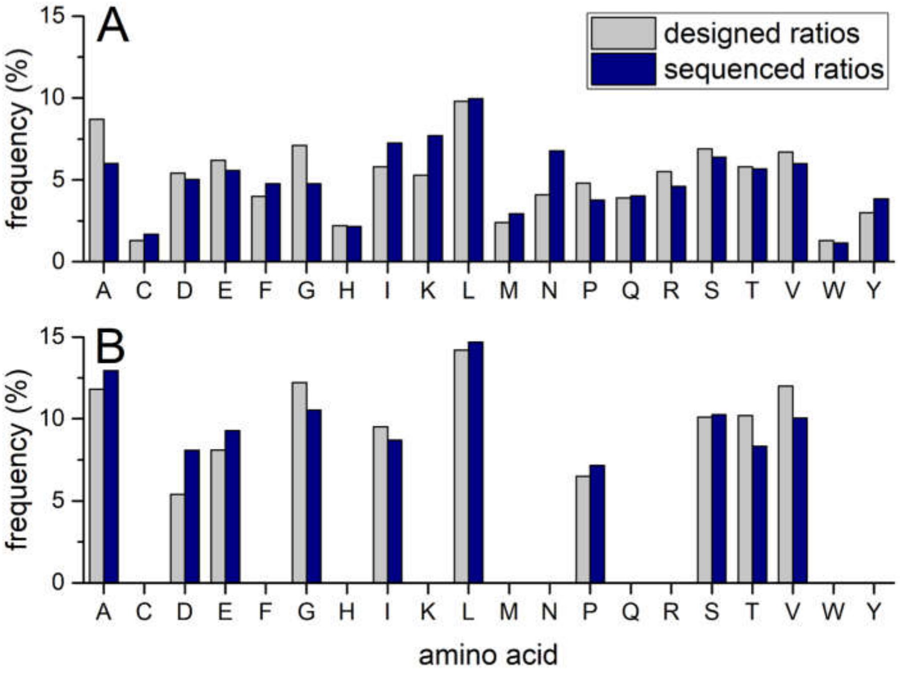


*Supporting Figure S3. Comparison of designed (grey) and experimental (blue) amino acid ratios in full (A) and early (B) alphabet libraries. Experimental amino acid distributions were calculated from the sequenced libraries DNA templates*

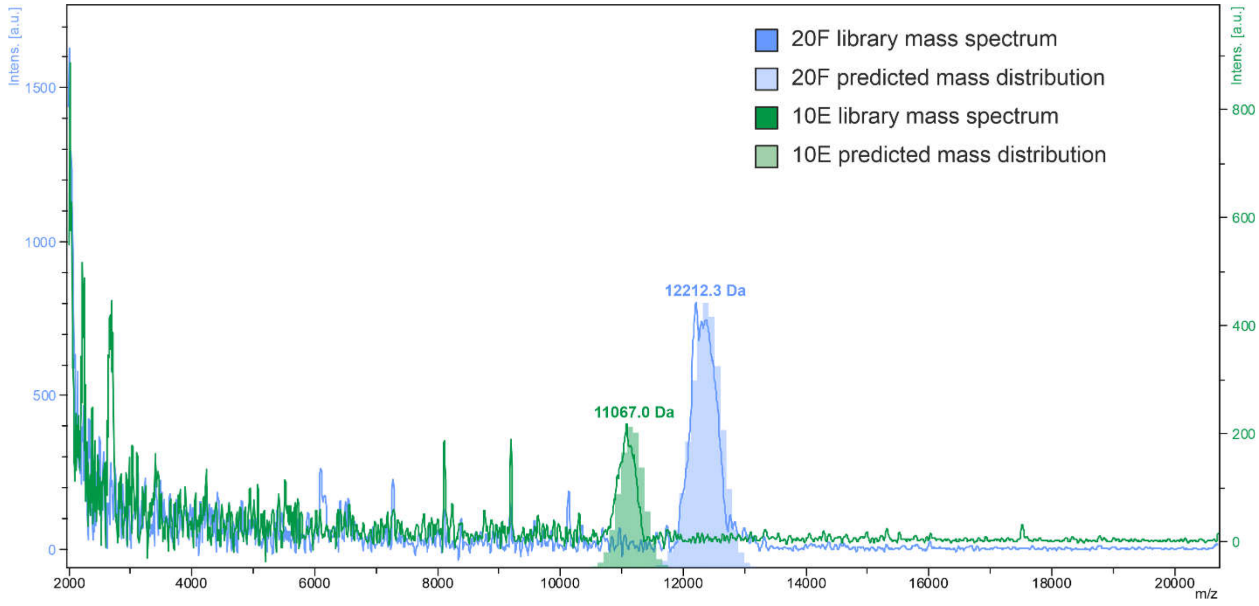

*Supporting Figure S4. Mass spectrometric analysis of purified full (blue) and early (green) alphabet libraries with their corresponding molecular weight distributions calculated in silico from the sequenced DNA templates*

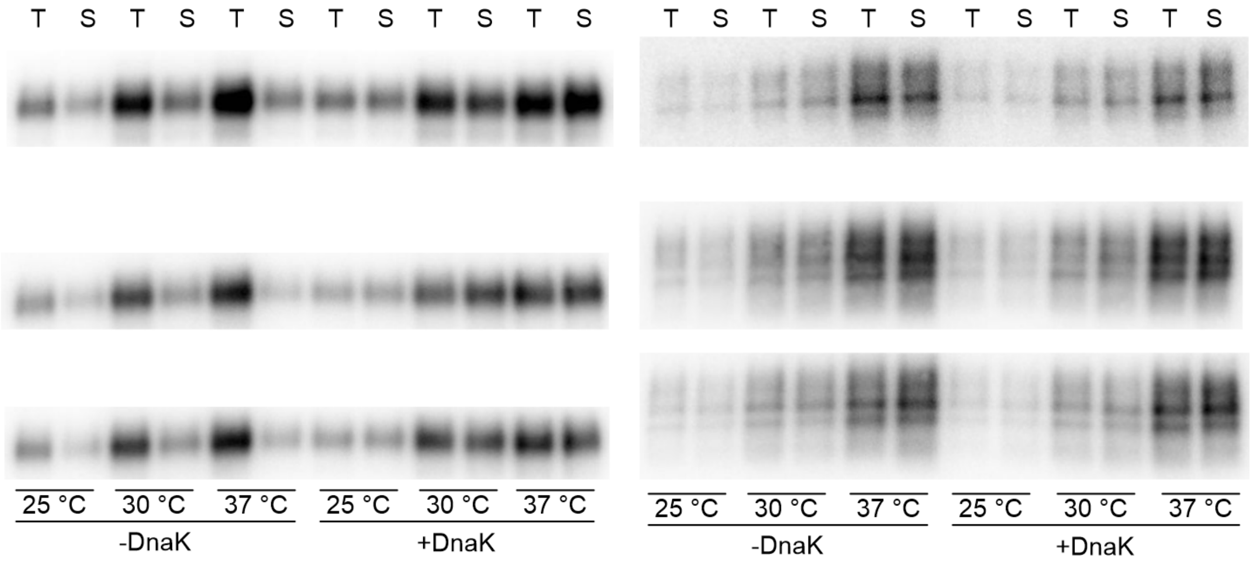


*Supporting Figure S5. Western blot signals used for temperature/chaperone dependent solubility analysis of full (A) and early (B) alphabet libraries. Total expressions (T) and soluble fractions (S) were analyzed*

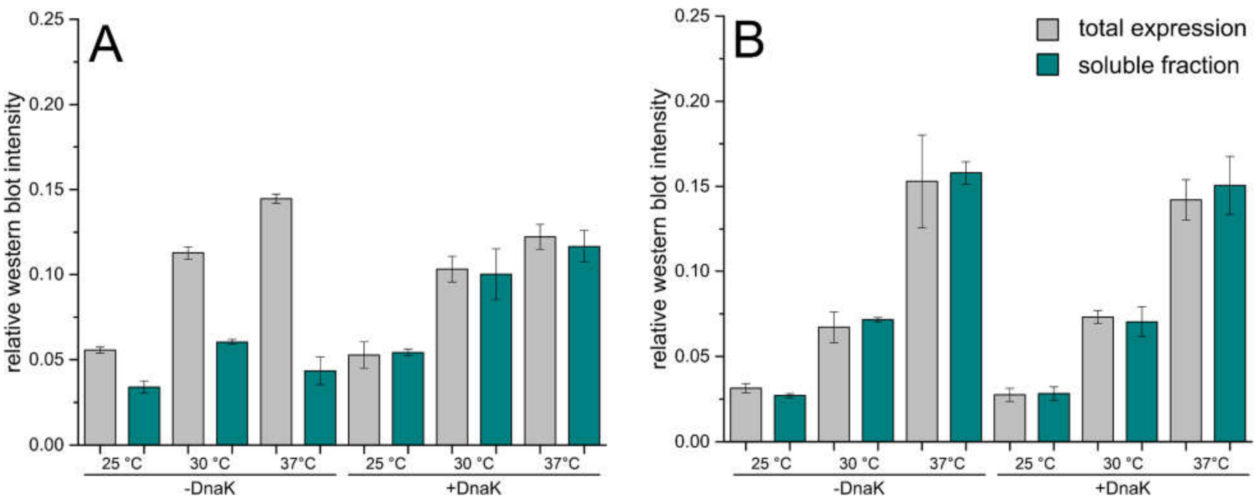


*Supporting Figure S6. Summary of triplicate western blot signal quantification of temperature/chaperone dependent solubility analysis of full (A) and early (B) alphabet libraries. Total expressions (grey) and soluble fractions (green) were analyzed*

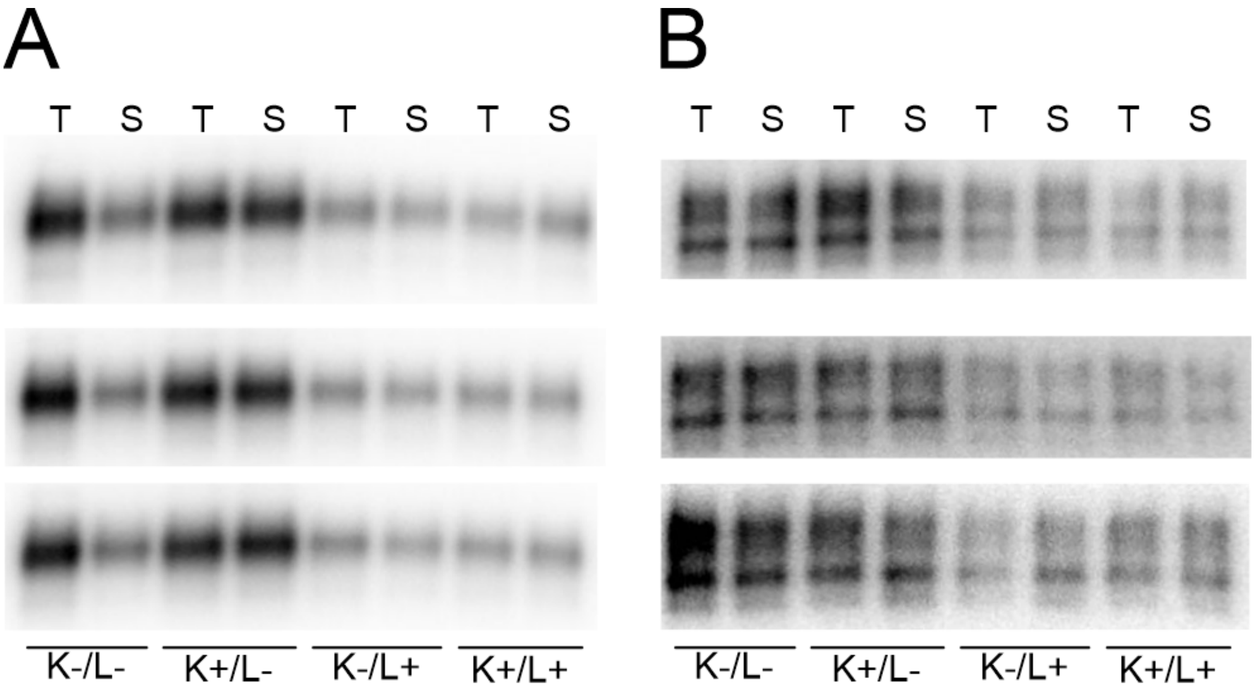


*Supporting Figure S7. Western blot signals used for Lon protease digestion/solubility analysis (K-/L-, K+/L-, K-/L+, K+/L+) and for the temperature dependent aggregation assay (42 °C) of full (A) and early (B) alphabet libraries. Libraries were expressed either in absence/presence of Lon protease (L-/L+) and absence/presence of DnaK/DnaJ/GrpE chaperone system (K-/K+). Total expressions (T) and soluble fractions (S) were analyzed. Only soluble fractions of chaperone absent (-K) or chaperone (+K) were analyzed after 42 °C heat shock treatment*

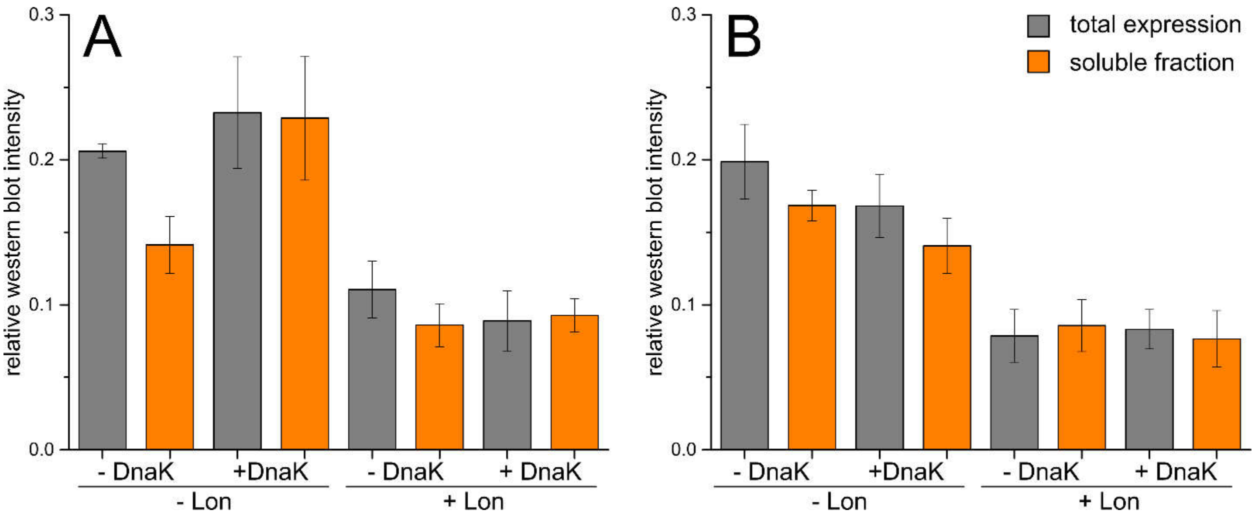

*Supporting Figure S8. Summary of triplicate western blot signal quantification of Lon protease digestion/solubility analysis of full (A) and early (B) alphabet libraries. Total expressions (grey) and soluble fractions (orange) were analyzed*

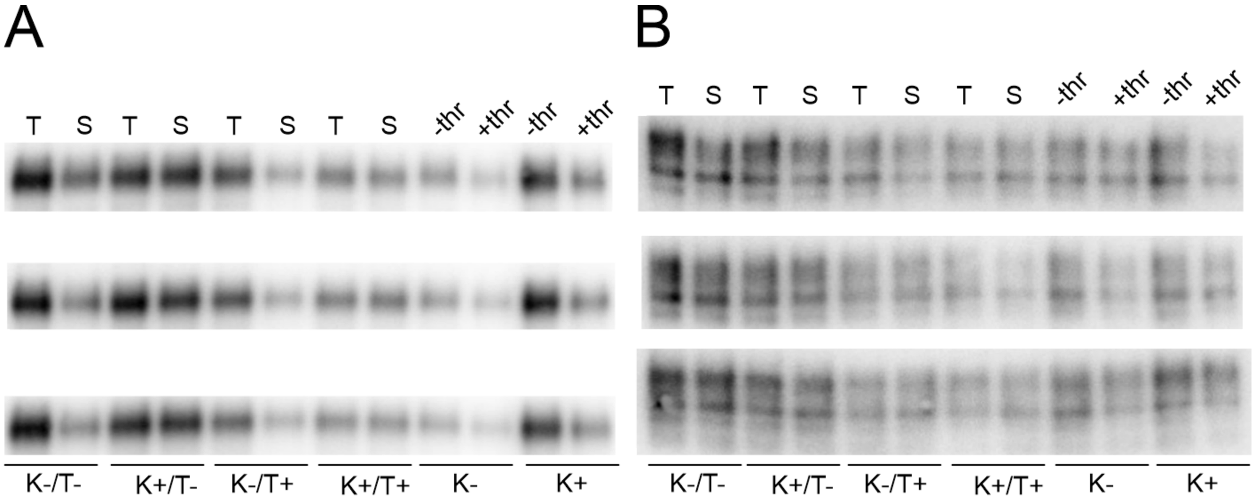


*Supplementary Figure S9. Western blot signals used for thrombin protease digestion/solubility analysis (K-/T-, K+/T-, K-/T+, K+/T+) of full (A) and early (B) alphabet libraries. Libraries were expressed either in absence/presence of the DnaK/DnaJ/GrpE chaperone system (K-/K+) and treated/untreated (T+/T-) by thrombin protease. Total expressions (T) and soluble fractions (S) were analyzed.*

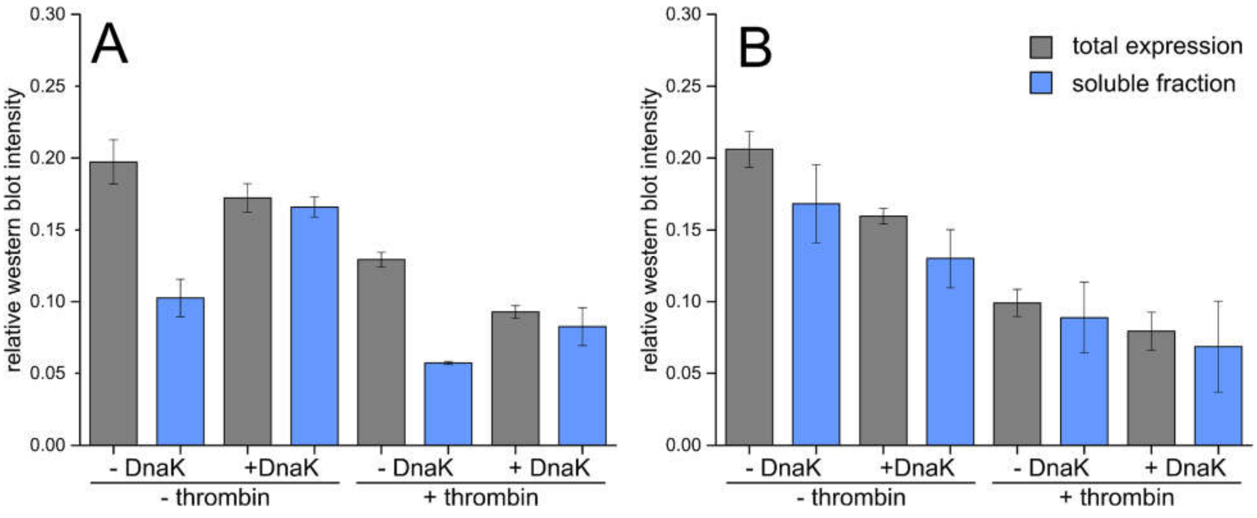

*Supporting Figure S10. Summary of triplicate western blot signal quantification of thrombin protease digestion/solubility analysis of full (A) and early (B) alphabet libraries. Total expressions (grey) and soluble fractions (blue) were analyzed*

**Supporting sequences**

>20F_full

CTGTAATACGACTCACTATAGGGACACCAATAGAGAAAGAGGAGAAATACTAGATGGATTA
TAAAGATGATGATGATAAGKHYKHYRMKDGKWTNNYARRSSHRVDGRMKRRSKHYMWMD
GKNYAGRRRMKNYAVDGRMKNYASMWSMWMWMKHYKHYVDGVDGKHYNYAMWMNYA
GCGTTAGTCCCGCGTGGGAGCNYASMWRMKKHYKHYVDGNYARMKNYAKHYVDGRRSVD
GKGSHHYNYAKHYRRSWTNVDGHHYRRSHHYHHYRMKNYANYAHHYSHRVDGHHYKHYD
GKVDGMWMWTNHHYRMKVDGKGSNYAVDGMWMNYAVDGRRSRRSSHRVDGRRSSMWRM
KCACCACCACCACCACCACTAA

>10E_full

CTGTAATACGACTCACTATAGGGACACCAATAGAGAAAGAGGAGAAATACTAGATGGATTA
TAAAGATGATGATGATAAGGNTHYAGNGHYAGNGKCWGNGGNGHYAHYAHYAHYAHYAG
NTHYAHYAHYAHYAHYAKCWGNGHYAGNTGAHHYAGNGHYAGNGHYAHYAHYAGNGGCG
TTAGTCCCGCGTGGGAGCGNTHYAGNGGNTGNGKCWGNGRBTHYAHYAGNGRBTHYAHYA
GNTHYAGNGHYAGNTHYAGNGRBTGNGGNGGNGRBTHYAHYAGAHHYAGNTGAHHYAHY
AGAHHYARBTGAHGNGHYAHYAGNTRBTHYAGNGGAHHYAGNGGMDGNTHYAGNGCACC
ACCACCACCACCACTAA
